# Generic principles of space compartmentalization in protocell patterns

**DOI:** 10.1101/2022.01.17.476586

**Authors:** Pierre-Yves Gires, Mithun Thampi, Sebastian W. Krauss, Matthias Weiss

## Abstract

Self-organization of cells into higher-order structures is key for multicellular organisms, e.g. during embryonic epithelium formation via repetitive replication of template-like founder cells. Yet, very similar spatial arrangements of cell-like compartments (’protocells’) are also seen in cell extracts in the absence of template structures and genetic material. Here we show that protocell patterns are highly organized, featuring a spatial arrangement and coarsening like two-dimensional foams but without signatures of disordered hyperuniformity. These features even remain unaffected when enforcing smaller protocells by stabilizing microtubule filaments. Comparing our data to generic models, we conclude that protocell patterns emerge by simultanous formation of randomly placed seeds that grow at a uniform rate until fusion of adjacent protocells drives coarsening. The strong similarity of our observations to the recently reported organization of epithelial monolayers suggests common generic principles for space allocation in living matter.

---

A hallmark of living matter is the ability to form and replicate well-defined cellular entities that self-organize into higher-order structures. Supposedly the most prominent example is the emergence of spatially organized tissues during the embryonic development of multicellular organisms: Starting from a single fertilized oocyte as a priming and pre-existing cell template, successive division cycles and rearrangements, invoking specific cell-cell interactions, eventually yield an organized array of cells [1]. Strikingly, these arrays often feature rather uniform cell sizes and very regular spatial arrangements, e.g. hexagonal arrays in epithelia [2, 3].

In contrast to this process that is driven by inheritance and replication, a spontaneous de-novo formation of ordered arrays of cell-like compartments (named ‘protocells’ hereafter) was recently observed in cell extracts from unfertilized oocytes of the amphibian *Xenopus laevis* [4]: Within a time course of about 30 min, ordered arrays of protocells with typical radii of some 100 *μ*m emerged spontaneously even without any organizing template structures like chromatin/DNA, centrosomes, or engulfing membranes (cf. Fig. 1). Despite being arrested in interphase and protein synthesis being inhibited, the protocell patterns are strikingly similar to epithelial monolayers [5] and spatially ordered arrays in early embryos [6] or oocytes [7]. Protocell assembly and self-organization is a genuine non-equilibrium phenomenon that is disrupted by ATP/GTP depletion, or by blocking the microtubule cytoskeleton dynamics or dynein motors; blocking actomyosin or kinesin motors involved in mitotic spindles had little to no effect [4]. Despite necessitating microtubules and molecular motors, protocells are not mere aster-like bundles, though, as they accumulate, for example, mitochondria [4]. Due to a reduced complexity (as compared to embryos), the emergence of protocells in the absence of priming template structures is an ideal model system for elucidating generic principles of spatial organization and compartmentalization in living matter.

**FIG. 1:**
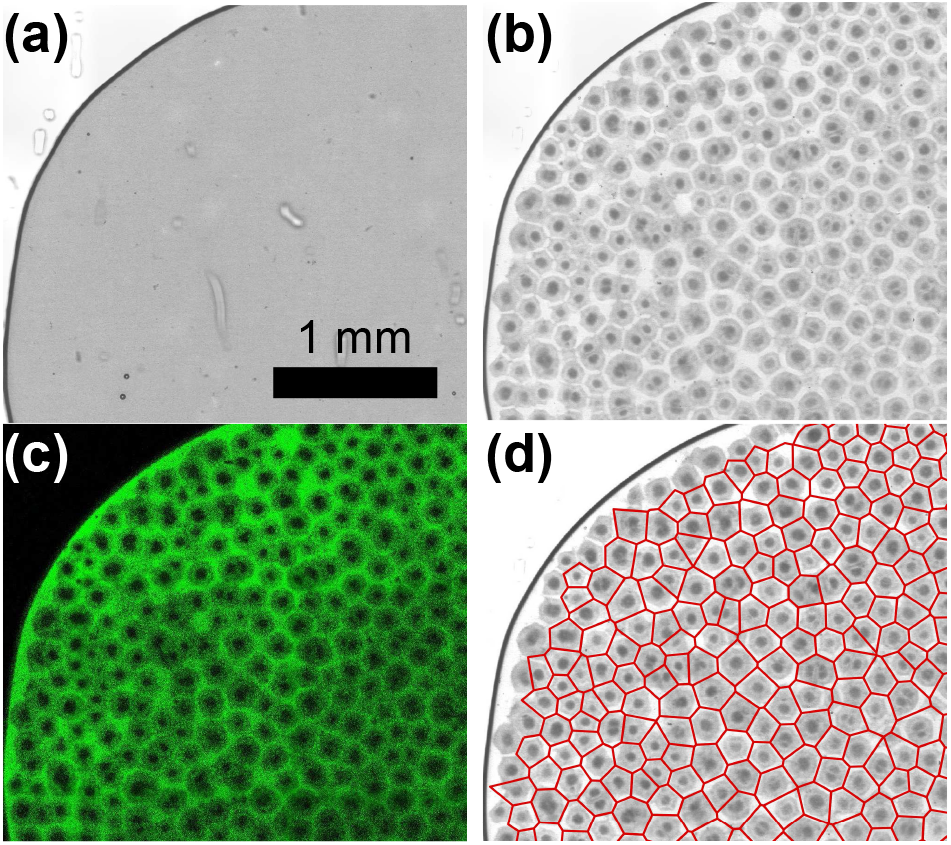
Cutout of representative bright-field images of an extract droplet (a) before and (b) after protocell formation. (c) Fluorescence imaging reveals that inert dextran molecules accumulate in boundary zones between protocells. (d) A Voronoi tesselation captures the essential geometry of the protocell pattern. See Fig. S4 for images of the complete droplet and movie 1 for its temporal evolution [8].

The fairly regular arrangement of protocells, akin to hexagonal arrays, suggests a particular organization that has attracted a lot of interest – disordered hyperuniformity [9]. Hyperuniform systems are characterized by a vanishing structure factor when the wave vector approaches zero, with disordered hyperuniform systems lacking the characteristic oscillations of strictly periodic systems. An alternative but equivalent criterion is obtained by considering the mean *μ* and the variance *σ*^2^ of the number of points found within a sphere of ra-dius *R*: For disordered hyperuniform systems, the normalized number variance Σ_2_(*R*) = *σ*^2^*/μ* approaches zero for *R* → ∞, indicating a long-range but non-crystalline order. In contrast, Poissonian random point patterns (PRPs) yield Σ_2_(*R*) = 1. Given the high degree of organization observed in epithelia and in protocell arrays, one may ask whether space allocation in biology strives for hyperuniformity.

Starting from this hypothesis, we have determined the geometric properties of protocell patterns and their coarsening dynamics over time. Following previous protocols (see Materials and Methods [8]), we were able to confirm the fairly robust emergence of protocell patterns in slab-like droplets of Xenopus extracts (see movie 1 [8]): Starting from homogenous droplets of freshly prepared extract (Fig. 1a), fairly regular arrays of protocell compartments emerged within 30-50 min after sealing the sample chamber (Fig. 1b). Notably, we did not observe the successive formation of individual protocell seeds at different spatial locations. Rather, the entire array became visible at some point (cf. movie 1 [8]), suggesting a global onset of pattern formation, akin to spinodal decomposition or Turing-pattern formation. After about 3 h, the pattern faded and started to disintegrate, most likely due to a lack of ATP/GTP molecules that are required for active processes.

Boundary lines between protocells had a lower absorbance in bright-field images, and hence a lower density than the interior of protocells. Most likely, radially organized microtubules are responsible for the increased crowding inside protocells as they constantly shuttle mitochondria and other organelles to the center region [4]. Exploiting and highlighting this permanent radial influx, we added minute amounts of accessory tracer beads (diameter 1 *μ*m) to freshly prepared extract droplets, that eventually accumulated in the center of protocells. The addition of these tracers was not seen to perturb proto-cell formation but enhanced the contrast for subsequent image analysis.

Unlike the rather large tracer beads, inert fluorescently labeled macromolecules (FITC-coupled dextran) were almost excluded from the interior of protocells and instead accumulated at the boundaries (Fig. 1c). This exclusion of inert macromolecules from densely crowded regions is similar to observations inside living culture cells [10]. Due to the good contrast between bright boundary zones and dark protocell centers, an automatic segmentation of bright-field images via a Voronoi tesselation was possible (cf. Fig. 1d and Figs. S3 and S4d [8]), facilitating a quantification of the protocells’ geometry over time.

As a first step, we analyzed the local geometry of proto-cells at two different time points, i.e. right after the first emergence of the pattern and 1-2 h later. To this end, we extracted individual protocell areas *A*, perimeters *L*, and vertex numbers *n*_*v*_ from images of different experiments and times. Accounting for varying average protocell areas, we determined normalized areas *A*_*n*_ = *A*/⟨*A*⟩ and perimeters *L*_*n*_ = *L*/⟨*L*⟩ by dividing out the mean values of the respective image. Since a Kolmogorov-Smirnov test (5% level) did not indicate significant differences of these normalized quantities between different experiments [11], we combined these for comparable time points into the same set and inspected their probability density functions (PDFs) *p*(*A*_*n*_), *p*(*L*_*n*_), and *p*(*n*_*v*_).

In line with the visual impression of a highly regular appearance of protocells, *p*(*n*_*v*_) highlighted a predominant occurrence of hexagonal protocells with appreciable probabilities also for pentagons and heptagons (Fig. 2a); polygons with more vertices were rare. This observation agrees well with epithelial monolayers [5] and aster patterns in *Phallusia* oocytes [7] but is in strong contrast to a Voronoi tesselation of PRPs for which the PDF is markedly wider (see Fig. 2a). The PDF of normalized areas, *p*(*A*_*n*_), assumed a narrow shape around a peak at unity, with a more than fourfold smaller variance than for a Voronoi tesselation of PRPs (Fig. 2b). The PDF of perimeters showed a very similar characteristics (Fig. 2b, inset), and both compare favorably to findings on epithelial cell layers on substrates of different rigidity [5]. Notably, all PDFs were independent of the time point at which they were acquired (cf. blue histograms vs. symbols in Fig. 2). This was also true for the PDF of protocell compactness (Fig. S5a [8]). These observations suggest rather uniform geometrical properties of proto-cells, e.g. being slightly disordered hexagons.

**FIG. 2:**
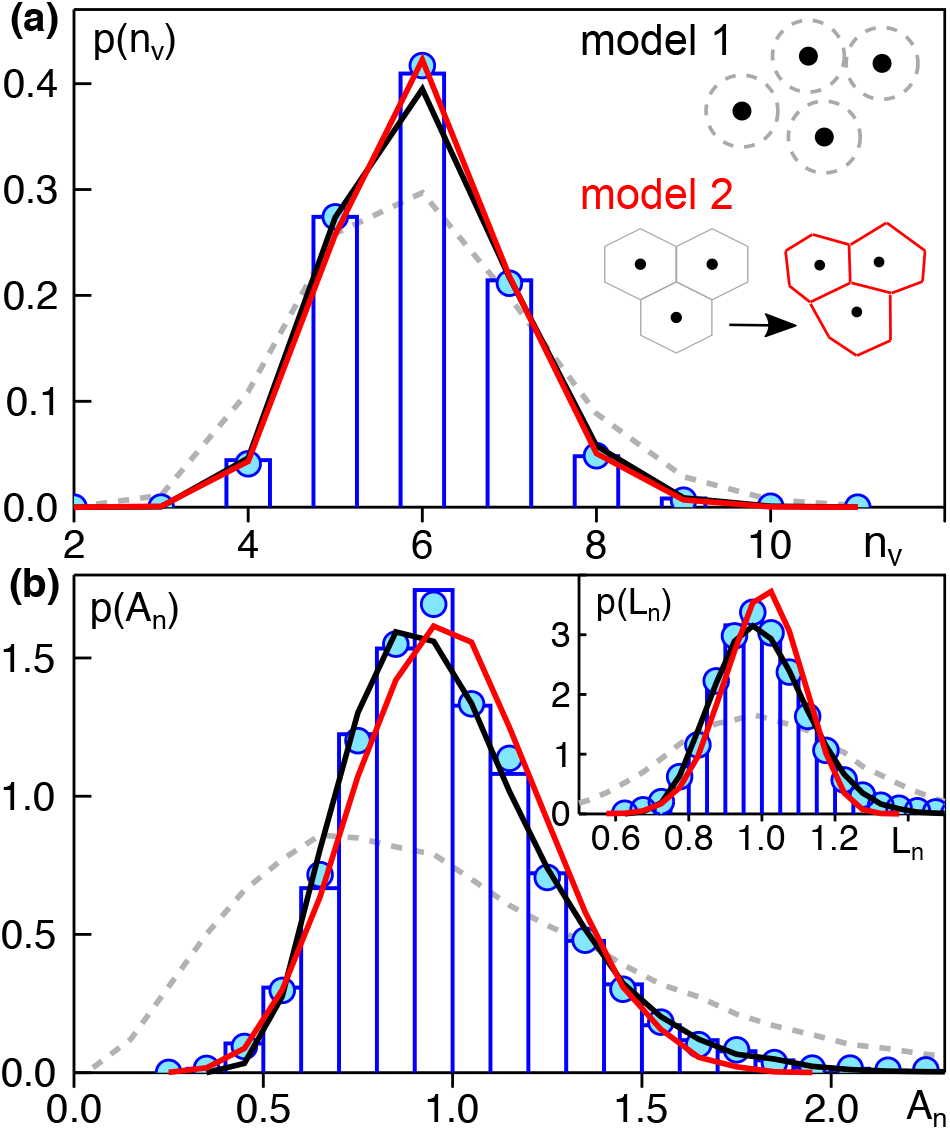
(a) The PDF of the vertex number, *p*(*n*_*v*_), of proto-cells right after the first emergence of the pattern (blue histogram) and 1-2 h later (blue circles) is very similar. Hexagonal cells are the most frequent phenotype, followed by appreciable amounts of pentagons and heptagons. The PDF for PRPs (grey dashed line) is markedly different. The experimental data are well captured by model 1 (*α*_1_ = 0.55, black line) and model 2 (*α*_2_ = 0.45, red line), both sketched in the inset and defined in the main text. (b) The PDFs of normalized cell areas, *p*(*A*_*n*_), and cell perimeters, *p*(*L*_*n*_), show a similar characteristics (color-code as before): Both models match the experimental data for early and late stages of the pattern, and the result for PRPs is markedly different.

Since protocell formation goes hand in hand with a focussing of microtubules into aster-like structures [4], we reasoned that stabilizing these cytoskeletal filaments may affect the protocell pattern. Since taxol had been described to stabilize microtubules without inducing major changes to cytoskeletal arrangements [12], we supplemented fresh extracts with this anti-cancer drug at different concentrations [8]. As a result, we observed that an increasing taxol concentration led to patterns with decreasing protocell sizes (Fig. S6 [8]). Quantifying the average protocell area ⟨*A*_80_⟩, found 70-90 min after starting the experiment, confirmed this visual impression, i.e. a considerable reduction of ⟨*A*_80_⟩ for increasing taxol concentration was observed (Fig. 3a). Similar observations have been reported for microtubule asters in oocytes [7].

**FIG. 3:**
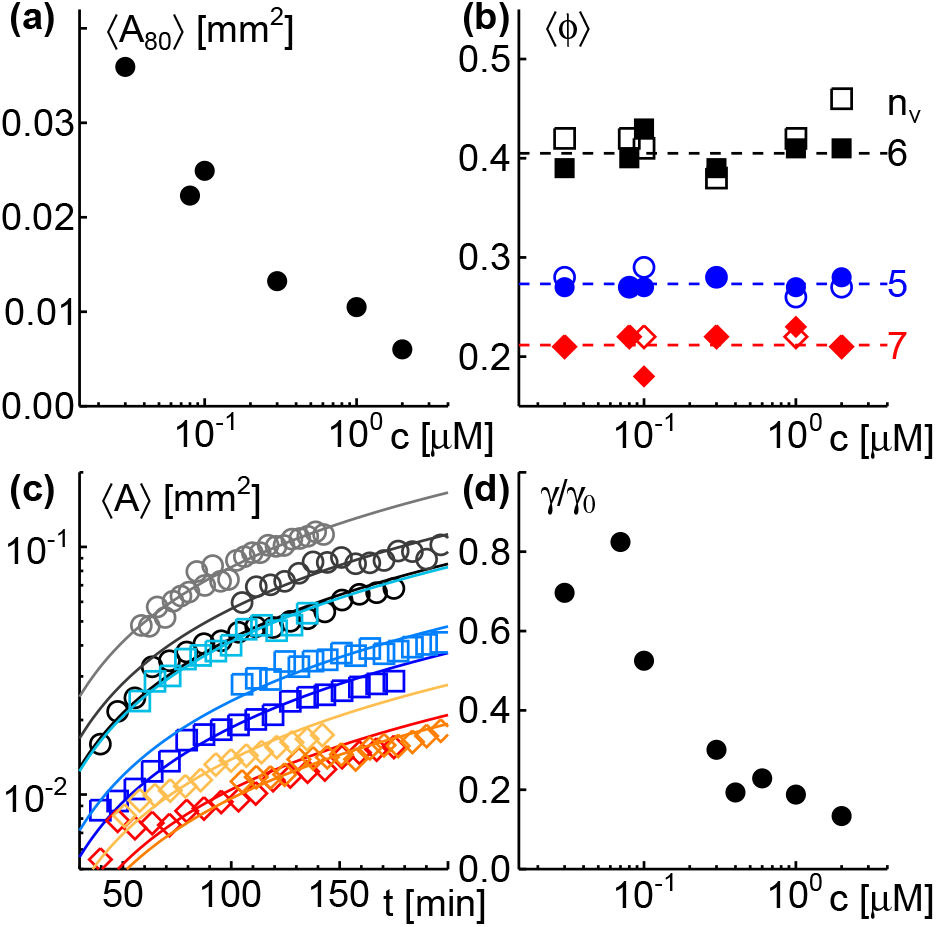
(a) The average protocell area ⟨*A*_80_⟩, found 70-90 min after starting the experiment, decreases for increasing taxol concentrations, *c*. (b) The average fraction ⟨*ϕ*⟩of pentagonal, hexagonal, and heptagonal protocells (blue circles, black squares, and red diamonds) is almost constant for all taxol concentrations *c*, irrespective of the time after the pattern emerged (open and filled symbols: right after pattern emergence and 1-2 h later). (c) Representative time courses of the average protocell area ⟨*A*⟩ for different taxol concentrations (grey circles, blue squares, red diamonds: *c* = 0, 0.1, 1 *μ*M) are well captured by a linear relation, ⟨*A*⟩ = *γt* (full lines); please note the logarithmic y-axis. (d) The growth rate *γ* decreases with increasing taxol concentrations, *c*, leveling off at about 15-20% of the rate observed for untreated extracts, *γ*_0_.

Quantifying *p*(*n*_*v*_), *p*(*A*_*n*_), and *p*(*L*_n_) in the presence of taxol did not reveal marked changes, irrespective of evaluating images right after the emergence of the pattern or 1-2 h later: The mean average fractions ⟨*ϕ*⟩ of pentagons, hexagons and heptagons was virtually unaltered (Fig. 3b), and the PDFs *p*(*A*_*n*_) and *p*(*L*_*n*_) for high taxol concentrations assumed the same shapes as the data shown in Fig. 2b (see Fig. S5b,c [8]). Thus, taxol treatment maintained all geometrical features of the pattern and only reduced the intrinsic length scale. The latter is most likely rooted in a reduced fraction of long microtubules in taxol-treated extracts [8].

To gain insights into the pattern dynamics, we monitored the average protocell area ⟨*A*⟩ as a function of time for varying taxol concentrations. In all cases, we observed a roughly linear growth ⟨*A*⟩ ≈ *γt* with a decreasing growth rate *γ* for increasing taxol concentrations (Fig. 3c,d). Notably, the number of protocells showed a decrease ∼ 1*/t* (Fig. S7a [8]), in accordance with the sum of all protocell areas being constant. Coarsening of the pattern was mainly due to a merging of protocells (see movie 2 and Fig. S7b for an example [8]).

Our experimental results therefore reveal that proto-cell patterns feature the same statistical scale invariance as two-dimensional foams [13]: The PDF of normalized areas is time-independent, irrespective of any taxol treatment, and the average protocell area grows linearly in time by a coarsening process at a conserved total area.

Given that two-dimensional foams can feature both, a long-range order with signatures of disordered hyperuniformity [14] but also a non-hyperuniform random patterns [15], we next probed protocell patterns on this aspect via the normalized number variance of protocell centers, Σ_2_(*R*). For hyperuniform systems, Σ_2_(*R*) should monotonously decrease for increasing test radii, *R*. The experimental data showed, however, a rapid and clear saturation at Σ_2_ ≈ 0.3, right after the onset of pattern formation and also 1-2 h later (Fig. 4). This indicates that protocell patterns are not hyperuniform at any time point, even though a visual inspection may suggest a near-crystalline order. Assuming values Σ_2_(*R*) < 1, the pattern can be viewed to have geometric properties of a hard-sphere fluid [9]. As a caveat, please note that only an asymptotic vanishing of Σ_2_(*R*) can properly reveal hyperuniformity, i.e. the finite sample size and potential inhomogeneities of protocell densities might mask a hyperuniform signature on the length scales available here.

**FIG. 4:**
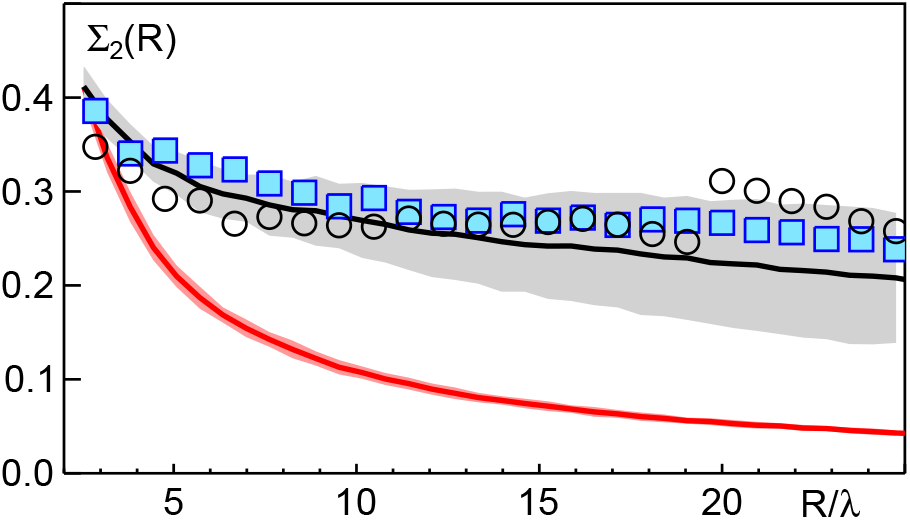
The normalized number variance Σ_2_ as a function of the rescaled test radius, *R/λ*, converges to a small but nonzero constant for the array of protocells (blue squares and black circles: time points right after pattern emergence and 1-2 h later, respectively). This indicates that the pattern displays no disordered hyperuniformity. While model 1 matches the experimental data well (black line) the hyperuniform characteristics of model 2 (red line) is clearly inconsistent with the experiment. Grey and red-shaded areas indicate the standard deviation for different realizations of the point patterns in the respective model.

To rationalize our experimental findings and to gain deeper insights into protocell self-organization, we formulated two simple models of how cell centers attain their spatial arrangement. Given that the pattern emerged spontaneously at some time, rather than a successive emergence of individual protocells at different loci, we assumed in both models an instantaneous existence of *N* center points that are placed on the unit square, hence defining a typical length scale 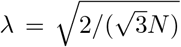 (chosen in accordance with [16]). Using a tunable parameter *α* ∈ [0, 0.7], two-dimensional point patterns were created in both models with the following rules (cf. sketches in Fig. 2a, inset):

1. Cell centers are chosen randomly from the unit square with the constraint that the minimal distance to neighboring centers is at least *αλ*.
2. Cell centers are created by displacing vertices of a triangular lattice by a distance *ξλ* in a random direction, with *ξ* ∈ [0, *α*] being a uniformly distributed random number.

Both schemes yield the steady-state pattern when cells emerge simultaneously, grow over time and stop growing upon contact. In model 1, random seed positions are combined with a uniform and isotropic growth rate, whereas model 2 assumes hexagonal cells (due to a global instability of the uniform state with a defined wave vector), perturbed by spatiotemporal fluctuations of the growth rate. Albeit not apparent immediately, the two models feature a very different long-range organization for *α* ≈ 0.5 (see Fig. S9 [8]): While model 1 displays a geometry that complies with a hard-disk fluid, indicated by Σ_2_ → *const*. < 1 for large radii [9], model 2 features an ever-decreasing number variance (Σ_2_ → 0) as expected for systems with disordered hyperuniformity [9].

Using *N* = 1000 points (for comparability with experiments), we produced *M* = 50 different realizations for each model and different choices of *α*. These point patterns were subjected to a Voronoi tesselation and evaluations of PDFs was done as for the experimental data. As a result, we observed that both models lead to a very good agreement with the experimental data when choosing *α*_1_ = 0.55 and *α*_2_ = 0.45 for model 1 and 2, respectively. In particular, PDFs for vertex number, cell area, and perimeter feature a shape that is empirically well captured by a gamma distribution [16], and all model PDFs match the experimental data so well that one cannot really claim a superiority of any model (Fig. 2). The PDF of the compactness, however, yields a first hint that model 1 might describe the experimental data somewhat better since model 2 predicts a slightly broader distribution (Fig. S5a [8]). Going beyond the local geometry, the PDF of next-neighbor correlations of protocells already provides evidence for model 1 being the more adequate description (Fig. S8 [8]). Finally, the normalized number variance Σ_2_ of model 1 captures the experimental data well, whereas the hyperuniform model 2 shows marked deviations (Fig. 4).

Based on our analysis, we can narrow down the emergence of protocell patterns to the following set of rules: Protocells emerge simultaneously at almost final positions, supposedly due to a low kinetic barrier for seed formation of local microtubule arrays. These seeds grow in an isotropic fashion at very similar growth rates by radial uptake of material from the close vicinity until touching neighboring protocells (growth stops). Adding taxol to stabilize microtubules only increases the number of seeds, leading to more but smaller protocells. Irrespective of tuning the length scale via taxol, the resulting pattern and its coarsening bear all features of a non-hyperuniform two-dimensional foam.

The stunning similarity of protocell patterns to epithelial cell monolayers [5] and aster arrays in *Phallusia* oocytes [7] suggests that this way of compartmentalization is a conserved feature in biological self-organization. Simulations of aster arrays [7] may even shed light on common self-organization events on microscopic length scales. It will therefore be interesting to explore quantitatively in more detail if other biological specimen, from cell monolayers to embryos, comply with the geometry of protocell patterns found here.

## Acknowledgments

Financial support by the VolkswagenStiftung (Az. 92738) and by the EliteNetwork of Bavaria (Study Program Biological Physics) are gratefully acknowledged. We thank A.C. Ramos and O. Stemmann (University of Bayreuth, Genetics) for providing Xenopus eggs, and A. Hanold for supporting the extract preparation.

## References

[1] B. Alberts, Molecular biology of the cell (Garland Science, Taylor and Francis Group, New York, NY, 2015), 6th ed., ISBN 9780815344643.

[2] A.-K. Classen, K. I. Anderson, E. Marois, and S. Eaton, Dev Cell 9, 805 (2005).

[3] K. Sugimura and S. Ishihara, Development 140, 4091 (2013).

[4] X. Cheng and J. E. Ferrell, Science 366, 631 (2019).

[5] S. Kaliman, M. Hubert, C. Wollnik, L. Nuic, D. Vurnek, S. Gehrer, J. Lovric, D. Dudziak, F. Rehfeldt, and A.-S. Smith, Phys. Rev. X in press, doi 10.1101/2021.04.10.439119 (2021).

[6] L. Guignard, U.-M. Fiúza, B. Leggio, J. Laussu, E. Faure, G. Michelin, K. Biasuz, L. Hufnagel, G. Malandain, C. Godin, et al., Science 369, eaar5663 (2020).

[7] N. Khetan, G. Pruliere, C. Hebras, J. Chenevert, and C. A. Athale, J Cell Sci 134, jcs257543 (2021).

[8] See Supplemental Material [url].

[9] S. Torquato, Phys Rep 745, 1 (2018).

[10] C. Donth and M. Weiss, Phys. Rev. E 99, 052415 (2019).

[11] No significant differences were detected between any two experiments on untreated samples or extracts with low taxol concentrations (≤ 0.1 μM); these data were hence combined.

[12] F. Verde, J. M. Berrez, C. Antony, and E. Karsenti, J Cell Biol 112, 1177 (1991).

[13] A. Saint-Jalmes, Soft Matter 2, 836 (2006).

[14] J. Ricouvier, P. Tabeling, and P. Yazhgur, Proc Natl Acad Sci 116, 9202 (2019).

[15] A. T. Chieco and D. J. Durian, Phys Rev E 103, 062609 (2021).

[16] H. X. Zhu, S. M. Thorpe, and A. H. Windle, Phil Mag A 81, 2765 (2001).

