## supplement for "Generic principles of space compartmentalization in protocell patterns"

### Generic principles of space compartmentalization in protocell patterns – Supplementary information

#### Contents

**I. Materials and Methods** 2
A. List of important reagents 2
B. Interphase extract preparation 2
C. Sample preparation and imaging 3
D. Image analysis and pattern detection 3
E. Taxol-induced change of the fraction of long microtubules 4

**II. Supplementary figures** 5

**III. Supplementary movies** 7

**References** 7

---

*These authors contributed equally

### I. MATERIALS AND METHODS

#### A. List of important reagents

- Human chorionic gonadotropin (Ovogest, 1000 U/mL)
- Aprotinin (Roche Diagnostics GmbH, 5 mg/mL in Milli-Q water)
- Leupeptin (Sigma-Aldrich, 10 mg/mL in Milli-Q water)
- Cytochalasin B (Calbiochem, 10 mg/mL in DMSO)
- Cycloheximide (Calbiochem, 10 mg/mL in Milli-Q water)
- Taxol (Sigma-Aldrich, 10 mg/mL stock solution in DMSO)
- 1,4-Dithiothreitol (DTT; Roche Diagnostics, 154 mg per liter of ELB)
- 1 $\mu$m polystyrene beads (Micromod Partikeltechnologie, micromer, COOH surface, product code: 01-02-103). Stock solution prepared by an aqueous 10$\times$ dilution of the pellet obtained by centrifugation.
- Fluorescein isothiocyanate-dextran (FITC-Dextran 10kDa; Sigma-Adrich). Stock solution: 10% (by weight) dissolved in Milli-Q water.
- Rhodamine labelled tubulin from bovine brain (Cytoskeleton Inc.)
- Guanosine 5'-triphosphate sodium salt hydrate (GTP; Sigma-Aldrich, 100 mM in Milli-Q water)
- General tubulin buffer: 80 mM PIPES pH 6.9, 2 mM $MgCl_2$, 0.5 mM EGTA.
- Aquapel water repellent glass treatment pack
- Egg Lysis Buffer (ELB): 250 mM sucrose, 2.5 mM $MgCl_2$, 50 mM KCl, 10 mM HEPES, pH 7.7 with KOH, 1 mM DDT.
- Dejellying solution: 2.0% (w/v) L-Cysteine (free base; Sigma-Aldrich) in Milli-Q water, pH 7.8 with KOH.
- 25x Marc's Modified Ringer's (MMR): 2.5 M NaCl, 50 mM KCl, 25 mM $MgCl_2$, 50 mM $CaCl_2$, 2.5 mM EDTA, 125 mM HEPES, pH 7.8 with NaOH.
- Secure-Seal$^{TM}$ imaging spacers (Grace Bio-Labs, SS8X9, 8-9 mm diameter, 120 $\mu$m height)
- Fluorinated ethylene propylene adhesive film (FEP; thickness 50 $\mu$m, Holscot Europe)
- Microscope slides (Corning, plain, 25 $\times$ 75 mm, thickness 0.96-1.06 mm)
- Cover slips (Menzel Gläser, 24 $\times$ 60 mm, #1.5 thickness)

#### B. Interphase extract preparation

The interphase extract preparation protocol was adapted from Refs (Deming and Kornbluth, 2006) and (Sparks and Walter, 2018) with the following minor modifications: A single HCG injection 16-17 h before the experiment was sufficient to obtain proper egg harvest. We changed the concentration of Marc's Modified Ringer's (MMR) to 0.5x (instead of 0.25x) and washed five times instead of three times. Correcting for a typo, we used KCl instead of HCl in the egg lysis buffer (ELB) preparation. Cycloheximide was added to the extract only after egg crushing (not in the ELB) but cytochalasin B was added already to the ELB in the centrifuge tube to which eggs were transfered for crushing (50 $\mu$g/mL final concentration).
Interphase-arrested cytoplasmic extracts were prepared from freshly laid oocytes of *Xenopus laevis* following standard protocols (Deming and Kornbluth, 2006; Sparks and Walter, 2018). In brief, eggs in the metaphase stage of meiosis II were collected and dejellied. These eggs were washed first with 0.5x MMR, then with ELB containing DTT, and finally packed by centrifugation with the excess buffer being removed. Packed eggs were crushed and fractionated into three distinctive layers by centrifugation using a Beckman Coulter JS-13.1 swinging bucket rotor and open-top polyclear centrifuge tubes (Seton, 4/6.5 mL). Then the mid cytoplasmic layer was carefully isolated and supplemented with 5 $\mu$g/mL of aprotinin, 5 $\mu$g/mL of leupeptin, 5 $\mu$g/mL of cytochalasin B, 50 $\mu$g/mL of cycloheximide, and stored on ice; these extracts were used within 6 h. The addition of cycloheximide inhibits protein synthesis, including the synthesis of new cyclin. Thus, the extract is arrested in an interphase state. The addition of protease inhibitors (aprotinin, leupeptin) limits protein degradation (Chan and Forbes, 2006), while cytochalasin B suppresses actin polymerization (MacLean-Fletcher, 1980). Some extracts were also supplemented with taxol to enhance microtubule stabilization without inducing major changes to cytoskeletal arrangements (Verde *et al.*, 1991).

##### C. Sample preparation and imaging

For imaging, interphase extract was supplemented with 1 $\mu$m polystyrene beads (1% by volume of stock solution) and FITC-Dextran (1% by volume of stock solution). Not adding any of these resulted in the same pattern formation but the contrast of protocell centers to the periphery was considerably dimmer. DMSO concentrations of all extracts were maintained at 1% by volume, if needed by compensating with pure DMSO. Extracts were mixed by gentle flickering, pipetting 3-4 times with a cut-off tip and carefully inverting the tube.
Fluorinated ethylene propylene (FEP) tapes were used to cover the bottom microscope slide and top glass cover slip that were separated by a 120 $\mu$m double-side tape spacer with circular 9 mm punched holes for hosting the extract droplets (see Fig. S1a); FEP coating was done one day before the experiment. To each of the holes, 4 $\mu$L of the extract was carefully pipetted into the center. Then, the whole chamber was sealed immediately with the coverslip to avoid evaporation. These samples were imaged with one of the following two microscopes in a tile-scan mode: a Leica DMI6000B inverted microscope with an HC PLS-APO 10x/0.30 DRY objective and a Leica DFC360FX camera or a Leica SP5 II confocal laser scanning microscope with an HCX PL APO CS 10x/0.40 DRY UV objective with 3.03 $\mu$m pixel width ($512 \times 512$ pixel tiles) and open pinhole. In both cases bright-field images were recorded, with an additional fluorescence channel on the SP5 for detecting FITC-Dextran (Excitation: 496 nm, Emission: 511-550 nm). The extract was imaged for more than four hours at room temperature (20°C) with a minimal time interval between consecutive tile scans.
For imaging of microtubules, rhodamine-labeled tubulin was first resuspended in 1 $\mu$L of 1 mM GTP-containing general tubulin buffer and 8 $\mu$L of fresh interphase extract. Samples for imaging were subsequently prepared by mixing 3 $\mu$L of this solution to 50 $\mu$L premixed interphase extract, hosted by a modified sample chamber for high-resolution imaging (Fig. S1b). Fluorescence imaging was performed with a Leica SP5 II confocal laser scanning microscope, using an excitation wavelength of 561 nm and a detection range of 575-650 nm. Images ($512 \times 512$ pixels, pixel size chosen according to Nyquist's theorem) were acquired with mono-directional scanning (scan frequency 400 Hz) within a maximum period of two hours, using a HCPL APO 100x/1.40 OIL or a HCXPL APO 63x/1.4 OIL CS2 objective.

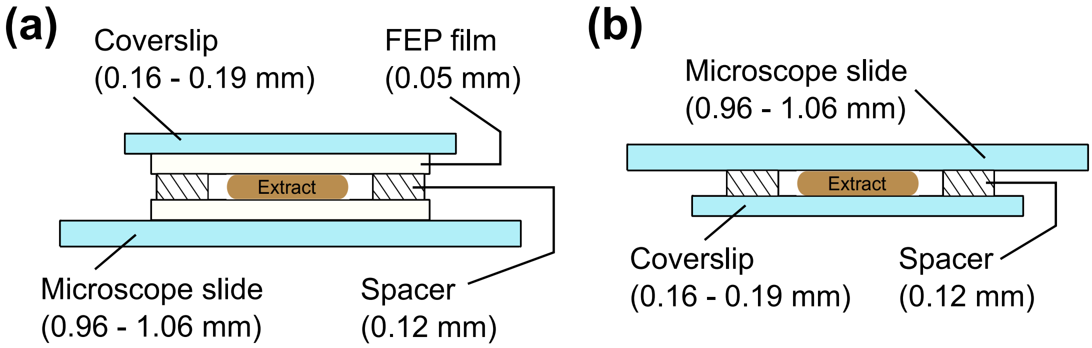


FIG. S1 Chamber design used for (a) imaging extract droplets and (b) fluorescence-imaging of microtubules (here, an aquapel-coating of the glass replaced the FEP film).

##### D. Image analysis and pattern detection

As a first step, shading and merging of images from a tile scan was performed with the Leica LAS X software using the options auto-stitching, smoothing, and linear blending, taking only the bright-field channel as a reference. Images from the Leica DMI6000B were resized by a factor 1/4 in each direction to arrive at a pixel dimension of 3 $\mu$m/px for all microscopes.
Using the time series of these reconstructed large-field images, the start and end times between which protocells were visible was determined visually for each droplet preparation. For these periods, the following operations were performed on the large-field images (see also Fig. S2 for illustration):

- Manual marking of the droplet boundary, defining a rectangular region of interest in each image for further processing.
- Automatic segmentation of the droplet boundary by detecting the ring of darkest pixels that is present at the droplet border.
- Binarization of the droplet image using the lowest of two thresholds that are obtained by Otsu's method (three-pixel classes with minimal intra-class variance, implemented via the *multithresh* function in Matlab).

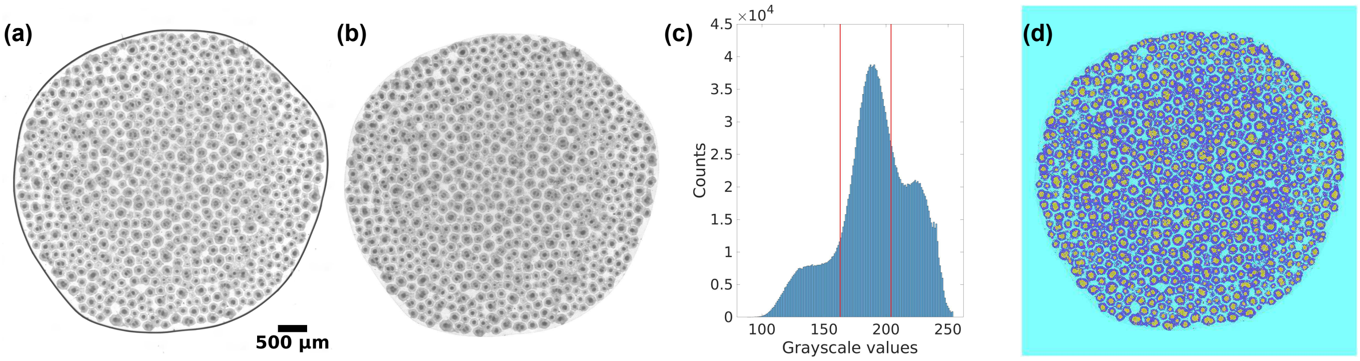


FIG. S2 (a) Representative example for an initial bright-field image of a droplet with protocell patterning. (b) Same droplet after masking (see text for details). (c) Pixel grayscale value histogram inside the droplet, with the two thresholds from Otsu's method indicated by vertical red lines. (d) Multi-thresholded droplet with darkest, intermediate, and brightest pixels assigned with colors yellow, dark blue, and light blue, respectively.

Subsequently, further filtering was performed to achieve a single-connected component of dark pixels within each protocell, defining the cell center. To this end, a sequence of classic morphological operations was applied (see Fig. S3 for illustration): Hole-filling (connectivity: four), image opening (using a $3 \times 3$ square), image dilation, hole-filling, and image erosion, with the same structural elements as before. Finally, connected sets of center pixels (highlighted in blue in Fig. S3d) were filtered according to their area histogram (using a bin width according to Scott's rule, connectivity for area: eight), i.e. the areas within the first histogram bin were rated as erratic pixels and hence removed (cf. examples before and after in Fig. S3d and Fig. S3e). The centroid positions of these filtered connected components were then used for a Voronoi tessellation (Fig. S3f).

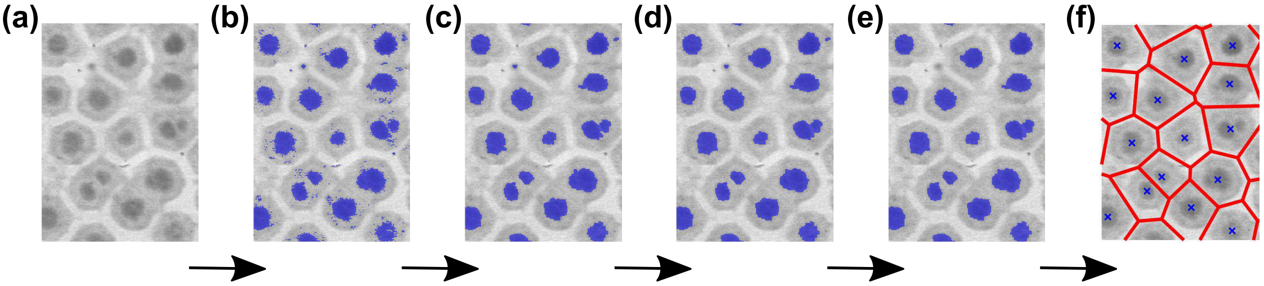


FIG. S3 (a) Close-up of a region with protocells in an extract droplet from bright-field imaging. (b) Same image thresholded according to Fig. S2c,d with the darkest pixels marked in blue. (c) Same after hole-filling and opening with a $3 \times 3$ square element to remove stray blue pixels. (d) After an additional image closing (same structural element) and intermediate hole-filling, protocell centers appear as single-connected blue area. (e) Filtering according to area histograms (see text) removes outliers. (f) Centroid positions of the obtained centers (blue crosses) support a meaningful Voronoi tesselation (red lines).

##### E. Taxol-induced change of the fraction of long microtubules

To extract microtubule lengths from fluorescence images, we used SOAX, a freely available analysis software for biopolymer networks (Xu *et al.*, 2015). Since very short filaments in semi-dilute and dense systems are difficult to assess with light microscopy without missing significant portions of this pool (unless fluorescent filaments are very sparse), we only considered microtubules with a length of at least 5 $\mu$m. From all detected microtubules of all images, we calculated the fraction of filaments with a length smaller than 20 $\mu$m, $f_s$, and the complementary fraction of longer filaments, $f_\ell$. The ratio $\zeta = f_s/f_\ell$ was taken as a self-normalizing measure for the frequency of long microtubules.
As a result, we observed that the average ratio $\zeta \approx 4.25$ in untreated extracts increased to an average of $\zeta \approx 9.54$ in extracts treated with 1 $\mu$M taxol. Thus, taxol treatment leads to a marked reduction of long microtubules in the extract. This result is in line with the effect of taxol as a drug that suppresses the dynamic instability of microtubules (Yvon *et al.*, 1999): In untreated extracts, the dynamical instability will lead to an outcompetition of small and shrinking microtubules by long and growing filaments, i.e. remaining microtubules grow longer on the expense of unsuccessful smaller ones that eventually vanish. In contrast, stabilizing all seeds by taxol increases the amount of growing filaments that compete for the same tubulin pool while growing with similar kinetics, eventually resulting in more but shorter filaments.

#### II. SUPPLEMENTARY FIGURES

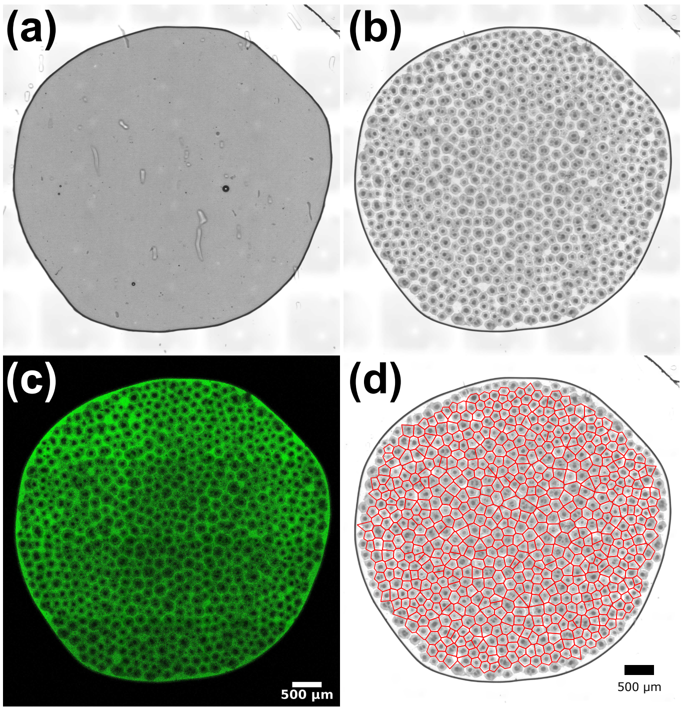


FIG. S4 Complete images of the cut-outs shown in Fig. 1 of the main text. (a,b) Bright-field images of an untreated extract droplet before and after protocell formation. (c) Fluorescence image of FITC-labeled dextran accumulating in boundary zones between protocells. (d) Voronoi tesselation of the protocell pattern. Images were taken (a) 7 min and (b,c,d) 175 min after chamber loading. The full temporal evolution is shown in `movie_1.avi`.

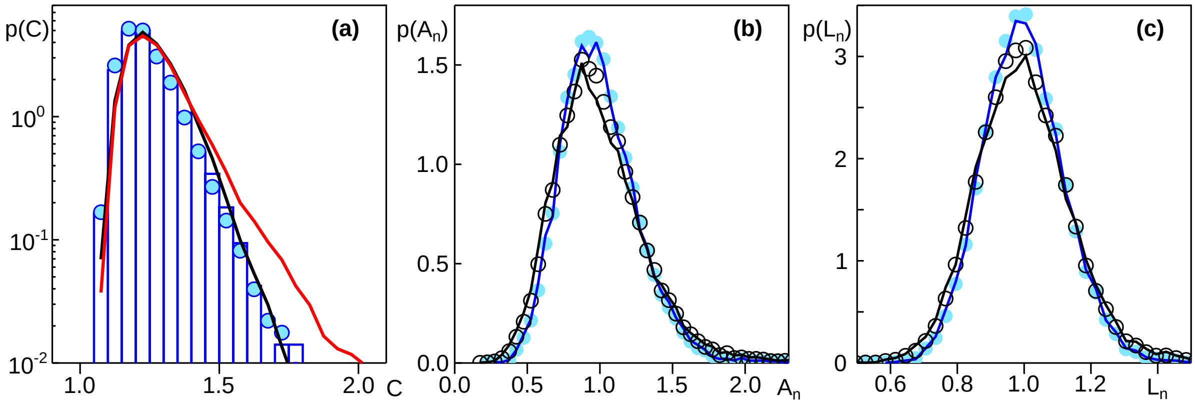


FIG. S5 (a) The PDF of protocell compactness, $p(C)$, with $C = \frac{L^2}{4\pi A}$, obtained for the same conditions as in Fig. 2 of the main text, features a mean $\langle C \rangle \approx 1.24$ that is larger than the value for circles ($C = 1$) and hexagons ($C = 6/(\pi\sqrt{3})$) but lower than that for squares ($C = 4/\pi$); color-code as in Fig. 2 of the main text. The experimental data are well captured by model 1 ($\alpha_1 = 0.55$, black line) and slightly less good by model 2 ($\alpha_2 = 0.45$, red line). Please note the semilogarithmic plot style. (b,c) The PDFs of normalized areas and perimeters for extracts that have been supplemented with 1 $\mu$M taxol or more (black lines and circles) agree well with the data for low and vanishing taxol concentrations ($c \leq 0.1$ $\mu$M, blue lines and symbols) right after the onset of pattern formation (lines) and 1-2 h later (symbols). A slight broadening of PDFs for high taxol concentrations most likely is caused by a decreased ratio of protocell size and space between protocells, resulting in an enhanced variability in the image analysis. NB: Data shown in blue are the same as in Fig. 2 of the main text with slightly smaller bin widths for an improved line visibility.

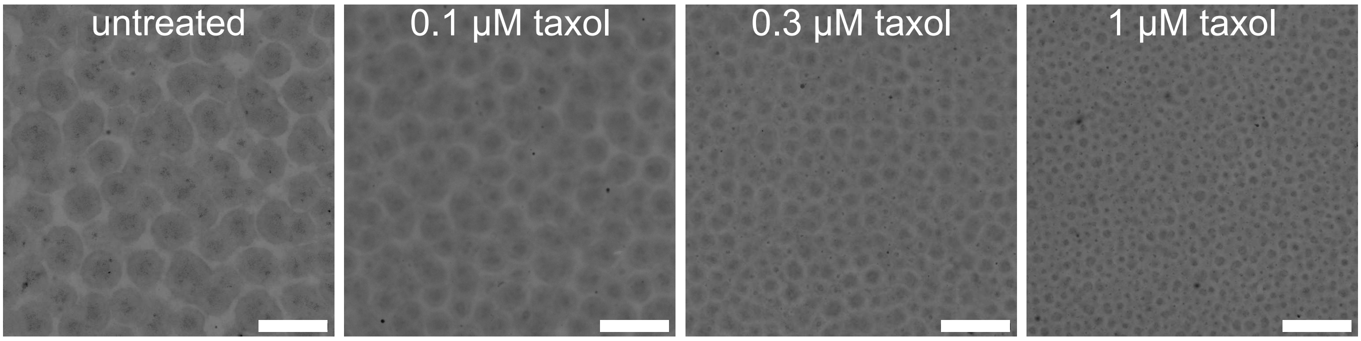


FIG. S6 Representative images of protocell patterns obtained from extracts without any treatment and different taxol treatments reveal a marked reduction of protocell sizes for increasing taxol concentrations; scale bars 100 $\mu$m.

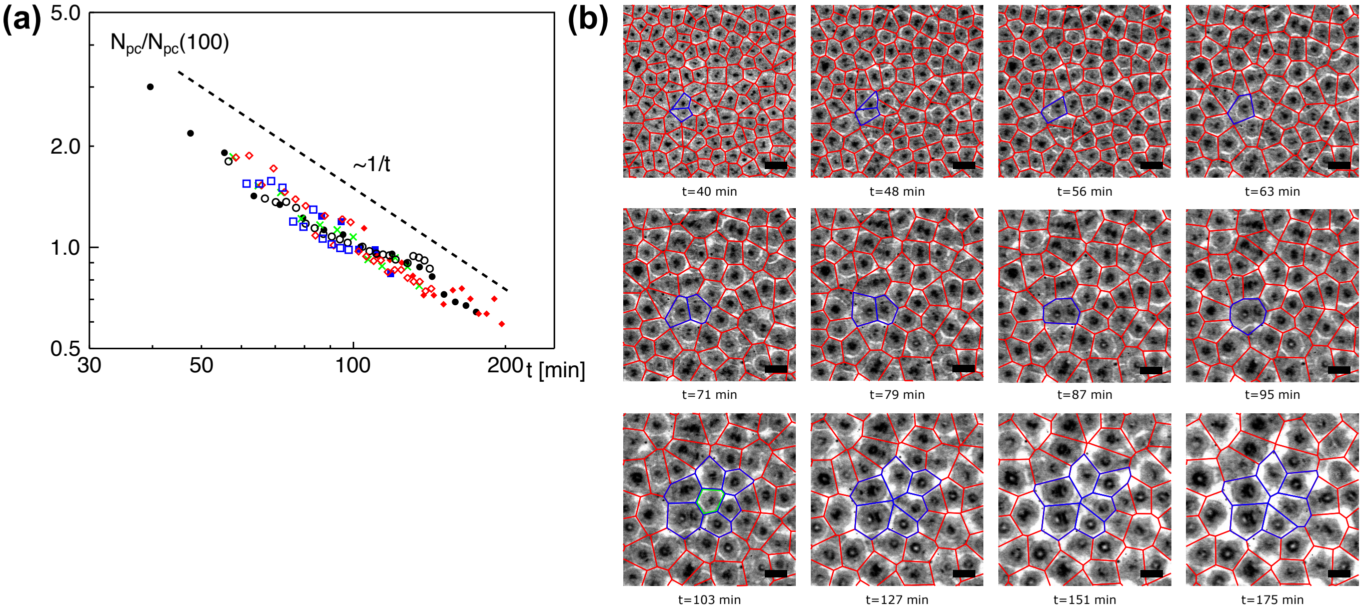


FIG. S7 (a) The number of protocells from different droplet preparations/experiments, normalized by the respective average protocell number at $t = 100$ min (different symbols), follow a common power-law decay $\sim 1/t$, hence compensating the linearly increasing average protocell area (Fig. 3c, main text) and securing a constant total area covered by protocells at all times. (b) Example for the successive coarse-graining of the pattern by merging of protocells; scale bars: 200 $\mu$m; see also `movie_2.avi` for the temporal evolution.

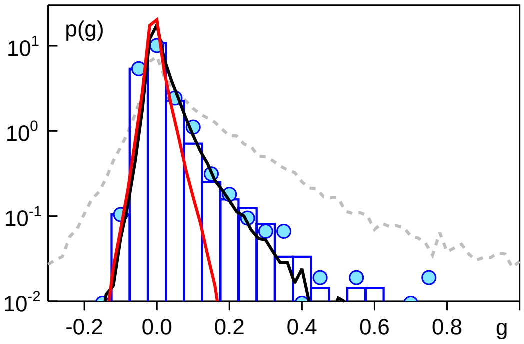


FIG. S8 The PDF of next-neighbor area correlation values, defined as $g_i = \left\langle (A_i - \langle A\rangle)(A_j - \langle A\rangle)/\langle A\rangle^2 \right\rangle_{j\in \mathrm{NN}(i)}$ for each cell with index $i$ and all its next-neighbor (NN) cells $j$ that share a common edge, is sharply peaked around zero for the experimental data right after the emergence of the protocell pattern (blue histogram) and after 1-2 h of coarse graining (blue circles). While model 1 captures the experimental PDF almost perfectly (black line), the hyperuniform model 2 decays too steeply for $g > 0$ (red line); the PDF for PRPs (grey dashed line) is far too broad. Hence, already on the scale of next-neighbor distances, the two models are clearly different, with model 1 being more consistent with the experimental data.

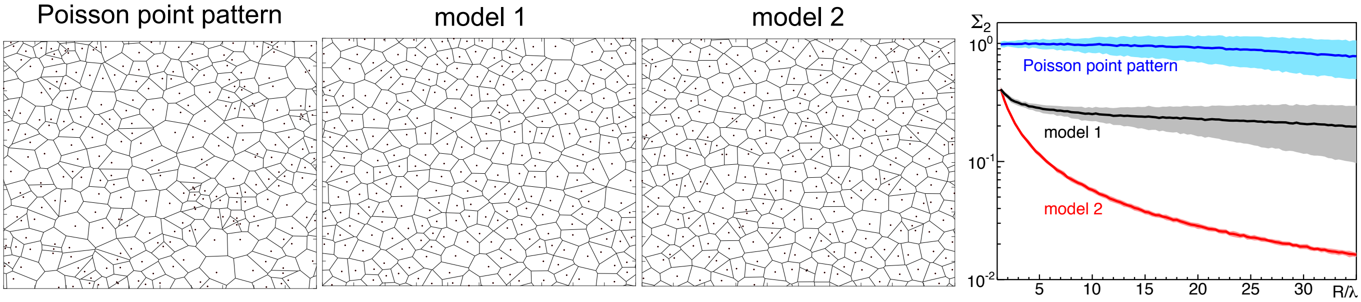


FIG. S9 Voronoi tesselation for a Poisson random point pattern, model 1 with $\alpha = 0.55$, and model 2 with $\alpha = 0.45$. The corresponding normalized number variance (shown in a semilogarithmic style) reveals that model 1 approaches a small but nonzero constant, whereas model 2 shows the typical feature of a disordered hyperuniform pattern, i.e. a decrease of $\Sigma_2$ towards zero; the Poissonian pattern shows the anticipated behavior $\Sigma_2 \approx 1$. Shaded areas indicate the standard deviation for different realizations of the point patterns.

#### III. SUPPLEMENTARY MOVIES

`movie_1.avi`: Temporal evolution of the extract droplet shown in Figs. 1 & S4 during protocell pattern formation.

`movie_2.avi`: Example for the temporal evolution of pattern coarse-graining shown in Fig. S7b .

##### References

Chan, R. C., and D. J. Forbes, 2006, in *Xenopus Protocols* (Humana Press), pp. 289–300.
Deming, P., and S. Kornbluth, 2006, in *Xenopus Protocols* (Humana Press), pp. 379–393.
MacLean-Fletcher, S., 1980, Cell **20**, 329.
Sparks, J., and J. C. Walter, 2018, Cold Spring Harbor Protocols **2019**(3), 194.
Verde, F., J. M. Berrez, C. Antony, and E. Karsenti, 1991, J Cell Biol **112**, 1177.
Xu, T., D. Vavylonis, F.-C. Tsai, G. Koenderink, W. Nie, E. Yusuf, I.-J. Lee, J.-Q. Wu, and X. Huang, 2015, Sci Rep **5**, 9081.
Yvon, A.-M. C., P. Wadsworth, and M. A. Jordan, 1999, Mol Biol Cell **10**, 947.
